# Neural mechanism underlying preview effects and masked priming effects in visual word processing

**DOI:** 10.1101/2023.07.24.550196

**Authors:** Xin Huang, Brian W. L. Wong, Hezul Tin-Yan Ng, Werner Sommer, Olaf Dimigen, Urs Maurer

## Abstract

Two classic experimental paradigms – masked repetition priming and the boundary paradigm – have played a pivotal role in understanding the process of visual word recognition. Traditionally, these paradigms have often been employed by different communities of researchers, with their own long-standing research traditions. Nevertheless, a review of the literature suggests that the brain-electric correlates of word processing established with both paradigms may show interesting similarities, in particular with regard to the location, timing, and direction of N1 and N250 effects. However, as of yet, no direct comparison has been undertaken between both paradigms. In the current study, we used combined eye-tracking/EEG to perform such a within-subject comparison using the same materials (single Chinese characters) as stimuli. Our results show the typical early repetition effects of N1 and N250 for both paradigms. However, repetition effects in N250 (i.e., a reduced negativity following identical-word primes/previews as compared to different-word primes/previews) were larger in the boundary paradigm than with masked priming. For N1 effects, repetition effects were similar across the two paradigms showing a larger N1 after repetitions as compared to alternations. Therefore, the results indicate that at the neural level, a briefly presented and masked foveal prime produces qualitatively similar facilitatory effects on visual word recognition as a parafoveal preview before a saccade, although such effects appear to be stronger in the latter case.

## 1. Introduction

In research on reading, two classic experimental paradigms – masked repetition priming (Forster et al., 1987) and the boundary paradigm (Rayner, 1975) – have played an important role in understanding the facilitatory effects of foveal and parafoveal information on word recognition. In the masked priming paradigm, a target word is preceded either by the same word or by an unrelated word as a briefly presented foveal prime. In the boundary paradigm, readers fixate target words for which either the same word or an unrelated preview had been shown in parafoveal vision during the preceding fixation. The facilitatory effects observed with both paradigms have been used to develop and refine models of reading; for example, the bi-modal interactive-activation model (Grainger & Holcomb, 2010) can explain masked priming effects, whereas the E-Z reader model (Reichle et al., 2006) can account for many of the effects of parafoveal previews on fixation times in reading. Yet, the results of both paradigms have been rarely directly related to each other, since behavioural studies with both paradigms have typically used different dependent variables (reaction times in masked priming paradigm and fixation durations in the boundary paradigm), making the effects difficult to compare. In contrast, electrophysiological recordings should allow for direct comparison between the facilitatory effects in paradigms. In the following, we briefly discuss both paradigms and then summarize the current study, which compared the effects of foveal and parafoveal facilitation in both paradigms directly within the same participants.

### 1.1. Masked repetition priming paradigm

In a typical masked priming paradigm, a foveal prime is presented for a brief duration (i.e., 40–72 ms) followed by a mask stimulus, which causes backward masking of the prime, making it consciously imperceptible. Behavioural responses (e.g., reaction times) to a subsequently presented target word are facilitated, if the target word is preceded by an identical word, even if there is a case change between prime and target (e.g., Forster et al., 1987; Forster & Veres, 1998). More recently, studies using event-related potentials (ERPs) have investigated the neural mechanisms involved in masked priming (Grainger et al., 2012; Holcomb et al., 2005; Holcomb & Grainger, 2006, 2007; Kiefer & Brendel, 2006). In alphabetic languages, (masked) repetition of visual words typically led to a reduction in two ERP components compared to unrelated words, the N250 and the N400 components (Chauncey et al., 2008; Holcomb & Grainger, 2006). The N250 repetition effect has a scalp distribution with the largest activation over midline and slightly anterior left hemisphere sites if it is recorded with a mastoid reference. At occipital electrodes, the effect shows a distribution with inverted polarity with a reduced negativity following repetitions, if an average reference is used (Huang et al., 2022). The N250 repetition effect seems to be sensitive to the visual-orthographic properties of the words, as it can be enhanced when primes and targets show orthographic overlap. In contrast, the N400 effect may reflect post-lexical semantic processing (see Kutas & Federmeier, 2011 for a review). In masked repetition priming studies, the N400 repetition effects shows a right-hemispheric lateralization (Chauncey et al., 2008; Holcomb & Grainger, 2006), with the largest activation over central-posterior scalp sites. The reduction of the N250 and N400 components may be considered as a correlate of the facilitation of orthographic and semantic processes (Grainger et al., 2012; Holcomb & Grainger, 2006).

In addition to the N250 and N400 effects, some studies also found repetition effects in the N1 component at occipital electrodes with an increased negativity for same-word versus different-word primes (Chauncey et al., 2008; Huang et al., 2022). This N1 component has also been called N/P150 component, as it is a highly focal bipolar ERP effect, with a positivity polarity at occipital scalp sites and a negativity polarity at anterior sites. Compared with the N250 and N400 effects, the N1 effect seems to be less consistent and less robust, possibly due to the use of mastoid references, a reference montage that is close to the electrodes where the N1 is maximal (Huang et al., 2022). In masked priming experiments, the early N1 component has been found to be not only sensitive to linguistic stimuli, like words (Holcomb & Grainger, 2006) and single letters (Petit et al., 2006), but also to non-linguistic stimuli, like pictures of objects (Eddy et al., 2007), suggesting that the N1 may reflect an early process involved in mapping visual features onto higher-level representations (Grainger & Holcomb, 2010).

Masked repetition priming studies in Chinese showed a similar pattern as alphabetic languages, that is, reduced N250 (Huang et al., 2022; Wong et al., 2014) and N400 amplitudes (Du et al., 2013; Wong et al., 2014; Zhang et al., 2012). These studies show that masked priming effects in N250 and N400 components are commonly obtained in visual word recognition irrespective of the properties of the writing system; hence, the N250 repetition effect might be a good candidate to compare facilitation effects at the level of early visual orthographic processing across different writing systems.

### 1.2. Boundary paradigm

In a boundary paradigm (Rayner, 1975), also known as the gaze-contingent display-change paradigm, readers read words normally with eye movements. When their eyes move from a pre-target word *n* to a target word *n* + 1, once the eye gaze crosses an invisible boundary in-between the words, the target word *n* + 1 is either replaced by another word or it remains the same word that was previewed in parafoveal vision during fixations on word *n*. In other words, fixations of word *n* + 1 occur either after a valid or after an invalid parafoveal preview of the word. The key finding in this paradigm is that fixation durations after valid previews are shorter than after invalid previews (the preview effect), suggesting that the response to a valid preview is facilitated compared to invalid previews (for a review, see Hyönä, 2012; Schotter et al., 2012a; Vasilev & Angele, 2017a).

At the electrophysiological level, studies combining EEG and eye-tracking have found a reduced negativity in fixation-related potentials (FRPs) following fixations on word *n*+1 if the preview was valid as compared to invalid (Dimigen et al., 2012; Kornrumpf et al., 2016). This effects, which has also been called “preview positivity”, is largest in a time window between 200 and 280 ms after fixating word *n* + 1 and maximal over occipital-temporal scalp sites. In subsequent studies, the preview positivity was not only observed in word list reading (Dimigen et al., 2012), but has also been replicated in reading natural sentences (e.g., Antúnez et al., 2022; Degno et al., 2019b, 2019a; Dimigen & Ehinger, 2021; N. Li, Wang, et al., 2022) as well as in studies using the RSVP-with-flanker-word presentation approach in Chinese readers (N. Li, Dimigen, et al., 2022; N. Li et al., 2015; N. Li, Wang, et al., 2022). While this effect has often been referred to as a “late N1 effect”, its time window and occipito-temporal scalp distribution are similar to those of the N250 component observed in masked priming studies (e.g., Holcomb & Grainger, 2007). Besides the preview positivity, a reduced N400 negativity to valid as compared to invalid preview has been observed between 300–500 ms after fixation onset (Dimigen et al., 2012; N. Li, Dimigen, et al., 2022), although not all studies show this effect (Degno et al., 2019b). In word lists, the N400 effects were only obtained with identical previews, whereas semantically-associated previews did not facilitate the process (Dimigen et al., 2012). However, in more natural reading setting (i.e., sentences), both identical and semantic associations seem to affect the fixation-related brain response (Antúnez et al., 2022; N. Li, Dimigen, et al., 2022; N. Li, Wang, et al., 2022), indicating that sentence meaning rapidly adapts to parafoveal preview.

Compared to the preview positivity and N400 effect, the *early* part of the N1 effect was less investigated or reported in previous studies (Dimigen et al., 2012; N. Li et al., 2015; Niefind & Dimigen, 2016). This N1 effect has a scalp distribution with the largest activation in the occipito-temporal regions, being more negative for valid than unrelated previews. According to the visual inspection on the waveforms of published studies, compared to the preview positivity, the N1 effect is usually smaller and less robust, and also less consistent.

### 1.3. Similarities and differences between masked priming and the boundary paradigm

Of course, the two paradigms differ in terms of the stimulation itself. First, the locations of primes/previews are different. As mentioned earlier, primes in masked repetition priming studies were usually fixated in the screen center (fovea), where participants passively read the words presented. In the boundary paradigms, previews usually appear in the parafovea of the right visual field, which projects information to the left hemisphere, therefore participants need to move their eyes actively. This means that there is a difference in attention orientation or attention allocation between the paradigms. A previous study comparing foveal-in-fovea priming to parafovea-on-fovea priming showed that foveal priming effects were bilateral but parafoveal priming effects were left-lateralized (Pernet et al., 2007), indicating that the locations of primes influence the neural processes. Therefore, it is possible that the foveal masked priming and parafoveal preview effect have different neural processes.

Second, in a masked repetition priming paradigm, the primes are usually masked with a backward mask (e.g., hash mark), which also causes a delay between primes and targets. In contrast, in a boundary paradigm, the preview is typically immediately replaced by the target, although this happens during the saccade towards the word (while there is strong saccade-related motion smear on the retina) (Schweitzer et al., 2023), which will mask some high spatial-frequency information. Therefore, it raises another question whether the existence of a delay and mask from prime/preview to target influences word recognition differently.

From a literature review, the N250 repetition effects appear to be similar in terms of time window and scalp distribution between masked priming and boundary paradigm. They both showed largest activation over occipito-temporal sites within a time window of around 180–280 ms. Previous studies (e.g., Dimigen et al., 2012; Holcomb & Grainger, 2006) have found that valid primes/previews could facilitate the neural process compared to invalid primes/previews, as reflected in a reduced N250 negativity for valid primes/previews compared to invalid primes/previews.

In terms of N400 effects, the two paradigms also seemingly show some similarities. In a time window of 300–500 ms, valid primes/previews elicited smaller amplitudes than invalid primes/previews. In masked priming studies, N400 effects usually have a posterior and left-lateralized maximum in an early phase but a more posterior and right-lateralized maximum at a later phase (Holcomb et al., 2005; Kiyonaga et al., 2007). In the boundary paradigm, the N400 effects usually have a centroparietal scalp distribution (like in masked priming studies), but they are less consistent, with highly robust N400 effects mainly found in RSVP-with-flanker paradigms (e.g., N. Li, Dimigen, et al., 2022; N. Li et al., 2015). In FRP studies, there evidence is more mixed; here, marginally significant effects (e.g., Dimigen et al., 2012), polarity-reversed effects (Degno et al., 2019a), but also robust N400 effects (N. Li, Wang, et al., 2022) have been reported.

With regard to the N1 component, a review of published figures suggests that the scalp distribution and direction of effects may be similar between masked priming and the boundary paradigm. However, given the fact that the early N1 repetition effect was rarely formally reported in previous boundary paradigm studies, it is not known whether the similarities generalize to all studies. The N1 repetition consist of an increased negativity for same versus different word presentations at occipital sites (Chauncey et al., 2008; Huang et al., 2022; Kornrumpf et al., 2016).

In terms of behavior measures, in masked priming studies, although the reaction times for repetition effects were not always reported (e.g., Chauncey et al., 2008; Holcomb & Grainger, 2007), the valid primes usually lead to shorter reaction times than unrelated primes (Morris et al., 2011; Morris & Stockall, 2012). In contrast, in boundary paradigms, fixation durations are usually analyzed, and it has been found that valid previews usually lead to 20-30 ms shorter fixation durations (for a review see Vasilev & Angele, 2017a).

Although the early N1 and N250 effects show apparent similarities, it is unclear, whether the early effects elicited in the two paradigms are actually the same. It is not known whether the same effects in terms of ERP amplitudes, topographies, and latencies are found if the same materials are presented in both paradigms. To establish better links between the literatures on both effects, it is important to understand the similarities and differences between the two paradigms using EEG measures. In the current study, for the first time, we used the same materials to investigate the neural correlates underlying the masked priming and boundary paradigm recorded in the same participants. We hope that this will also allow us and others to span a bridge between these two important lines of research and to cross-reference the results and their theoretical interpretations in the respective literatures. A direct comparison of the two paradigms may also allow us to understand the EEG components better, as previous discussions and functional interpretations typically only occurred within the framework of one paradigm (e.g., Dimigen et al., 2012; Grainger & Holcomb, 2009). Previously, in a to our knowledge unpublished master thesis, Pancani, 2016) compared the preview and priming effect with different groups of participants, but the masked repetition priming paradigm in her study used a blank screen as a forward and backward mask, which may make the prime consciously identifiable. In addition, her study focused more on later effects (N400 component), with little attention on early N1 repetition effect. Furthermore, she did not replicate the typical preview positivity when the previews and targets were presented in a different case; the authors argued that this was because of the salient visual change occurring in the unrelated and repeated condition disrupting reading. To rule out this possibility and to avoid visual overlap between primes/previews and targets, we therefore used different fonts even for the repeated condition.

### 1.4. Current study

Given what appears to be considerable similarities of ERP effects in masked priming and boundary paradigm studies on visual word recognition, it is important to directly compare these effects in the same study. Demonstrating a similarity of the neural facilitation effects that have long, but separate traditions in research on visual word processing would allow researchers to compare results better and link theoretical interpretations between the two lines of research. Therefore, in the current study, we compared neural facilitation effects between masked priming and boundary paradigms in the same participants, with the same stimuli, as well as the same EEG setup and preprocessing steps. We also used concurrent eye-tracking to co-register eye movements with the EEG.

All hypotheses and analyses were pre-registered at https://osf.io/hk37q/. Behaviourally, we expected to replicate the typical preview effects in terms of fixation durations in the boundary paradigm, although most studies obtained the preview benefit on eye movements through natural sentence reading (e.g., Yan et al., 2009; Yang et al., 2012) rather than during single character reading. Electrophysiologically, we expected that both paradigms would show an attenuated (less negative) N250 in response to a stimulus following an identical prime/ preview as compared to an unrelated prime/preview. This effect was expected to be larger for the boundary paradigm compared to the masked priming experiment based on previous studies (Dimigen et al., 2012; Petit et al., 2006), as active oculomotor behaviour seems to modulate word recognition under otherwise comparable conditions (Kornrumpf et al., 2016). With regard to the N1 component, the boundary paradigm was expected to result in an increased (more negative) early N1 after identical compared to unrelated previews. This effect was expected to be larger than in the masked priming experiment (i.e., interaction with factor *Paradigm*). The interaction might result from a reduced/absent effect in the masked priming experiment or even from an effect in the masked priming paradigm that reverses direction (larger negativity after related than unrelated stimuli; Grainger et al., 2012). For our analysis, we also used a time-point-to-time-point Topographic Analysis of Variance (TANOVA) that includes all electrodes to test at which time points the masked priming effect and preview effect would occur across all electrodes. As we were mainly interested in the early processes of visual word recognition, and since later components (e.g., P350 or N400) are much more susceptible to tasks (e.g., Huang et al., 2022), they were not included in our analysis.

## 2. Methods

### 2.1. Participants

Twenty-nine native Cantonese-speaking Chinese participants (13 males; mean age = 19.21 years, range = 18–25 years) were tested in both paradigms. Data from one additional participant was excluded from the analysis due to a small number of trials (average trial number in each condition < 15). All participants had normal or corrected-to-normal vision, with no history of dyslexia or ADHD (self-report). And they were all right-handed as determined by the Chinese Handedness Questionnaire (Li, 1983). Written informed consent was obtained prior to the experiment. All participants were reimbursed 50 Hong Kong dollars (about 7 USD) per hour. The study was approved by the Joint Chinese University of Hong KongLJNew Territories East Cluster Clinical Research Ethics Committee.

### 2.2. Materials

Eighty-four traditional Chinese characters were selected from the Chinese Character Database (Kwan et al., 2006) as target characters. These 84 characters were paired with either identical or unrelated characters, therefore, a total of 168 trials were included in the data analysis. The unrelated characters and target characters were matched according to stroke numbers (*t* _(166)_ = 0.04, *p* = 0.97), radical family size (*t* _(166)_ = 0.35, *p* = 0.73) and frequency (*t* _(166)_ = 0.07, *p* = 0.95). There were no shared radicals and homophones within a pair.

An additional 16 animal characters served as probes and were either presented in the prime/preview or target position; the probability of them being target vs. prime/preview character was equal (4%). These animal characters were once paired with unrelated characters and once paired with identical animal characters. In total, there were 100 pairs of trials with unrelated preview/prime and 100 pairs with an identical preview/prime. For trials containing animal names, participants were instructed to press a button (response hand counterbalanced across participants) whenever they detected an animal name in either the prime or target position. Trials with animal characters were excluded from the EEG data analysis. With this design, the stimuli used in the two paradigms were the same, including the characters indicating animacy. The order of items within each paradigm was randomized. The order of the two paradigms was counterbalanced across participants.

The animal names served as probe items in a semantic categorization task in which participants were instructed to rapidly press a single button whenever they detected an animal name in either the prime/preview or target position. A practice session was administered before the experiment to familiarize the participant with the procedure.

### 2.3. Procedure

#### 2.3.1. Masked priming paradigm

Visual stimuli were presented on a 24-in monitor (BenQ ZOWIE XL2411K, resolution: 1920×1080 pixels; vertical refresh rate: 144 Hz) and located at a distance of 90 cm directly in front of the participant. Stimuli were displayed in high contrast as black characters on a white background. As shown in Figure. 1B, each trial began with a fixation cross in the middle of the screen. Five hundred milliseconds later, a forward mask was presented for a duration of 500 ms. The forward mask was replaced at the same location on the screen by prime character (set in a Kaiti font) for 50 ms. The prime was then immediately replaced by a backward mask. The backward mask remained on screen for 20 ms and was immediately replaced by the visual target (in the PMingLiu font) for a duration of 500 ms. Target word presentation was followed by a 2000 ms long empty white screen. When the target disappeared, marking the end of the trial, participants were asked to press the button to indicate whether they have seen an animal name during the trial or not (regardless of the position of this probe as prime or target). In the masked priming paradigm, participants were asked to refrain from blinking and from moving their eyes while stimuli were presented.

**Figure 1.**
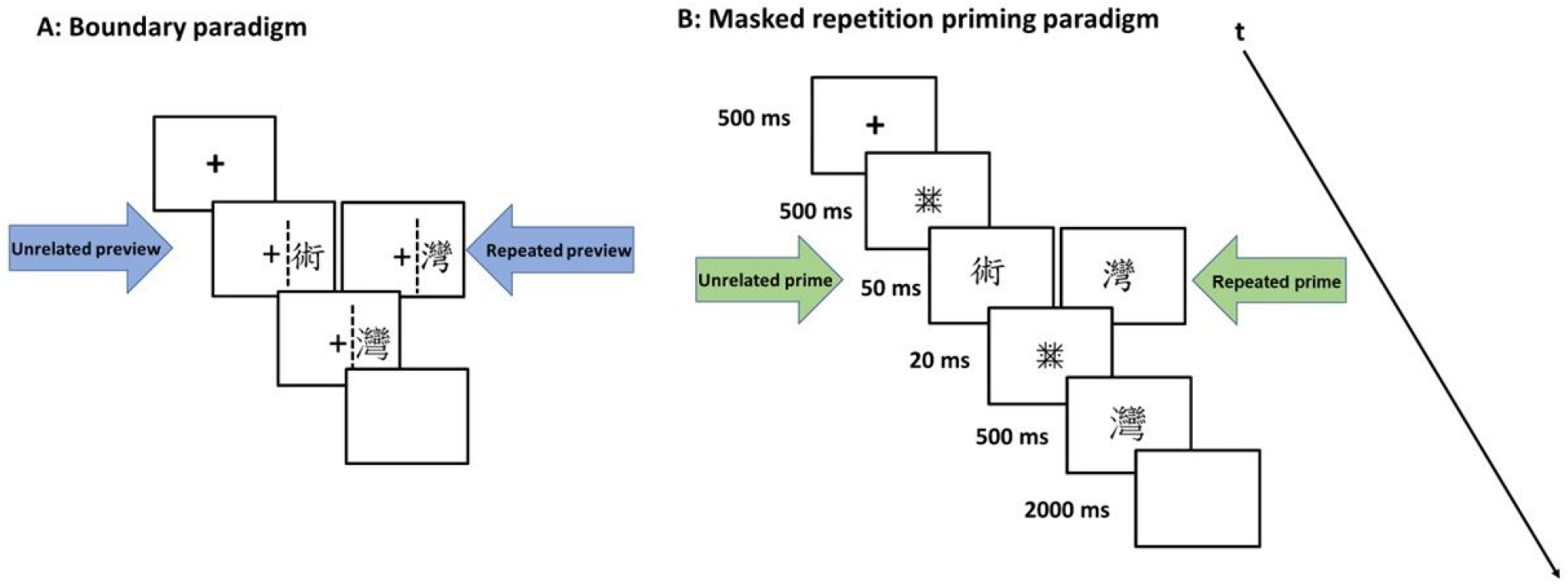
A schematic of a typical trial in the boundary paradigm (A) and the masked priming (B). The dashed line in the boundary paradigm indicated the invisible vertical boundary. Once the eyes cross this boundary, during the saccade, the preview is exchanged to the target. The figure shows an example of a trial in which the prime/preview was unrelated (left) and repeated/valid (right) to the target character, respectively.

#### 2.3.2. Boundary paradigm

Technical settings were the same as in the masked priming paradigm except for the schematic of the trial (see Figure 1A for a schematic of a typical trial). In the boundary paradigm, each trial started also with a fixation cross in the middle of the screen. Participants needed to fixate on this cross within 3 s (fixation check). Afterwards, the preview character appeared centered at a visual angle of 5.5° to the right of fixation, and participants moved their eyes rightwards towards it. The distance from the fixation cross to the left edge of the character was 3.03°. An invisible boundary was located in-between fixation cross and character, at an eccentricity of 2.5° to the right of fixation. The trial was aborted if the boundary was not crossed after 3000 ms. Analogous to the masked priming experiment, the preview character was presented in Kaiti font. Once the participants’ eyes moved across the invisible boundary, the preview was replaced by the target character in a PMingLiu font. Following the crossing of the invisible boundary, targets were presented for a duration of 1000 ms^1^. As in the masked priming experiment, once the target disappeared, there was a 2000 ms blank white screen and participants were then asked to press the button to indicate whether they had seen an animal name (again, regardless of the position of this probe as preview or target).

Display change awareness was assessed in a structured interview after both experiments. Participants were first asked whether they had noticed “anything strange about the visual display of the text” (White et al., 2005). If they answered “no”, they were informed that changes had taken place and asked again whether they had noticed any. If they did, participants were asked to estimate the number of changes perceived. Due to the simplified single-saccade setting of the current reading task, we expected that most participants would be aware of the display changes (from preview to target) in the boundary paradigm trials. It is also true for the masked priming paradigm as there were visual transient on the display. Notably, however, N1 effects in the boundary paradigm have been shown to arise regardless of display change awareness (Dimigen et al., 2012).

### 2.4. EEG recordings

The EEG was recorded from 64 Ag/AgCl scalp electrodes mounted in a textile cap at standard 10–10 positions and referenced online against CPz. Two electro-oculogram (EOG) electrodes were placed on the outer canthus of each eye and one EOG was placed on the infraorbital ridge of the left eye. Signals were amplified with an EEGO amplifier system (Advanced Neuro Technology, Enschede, Netherlands) at a bandpass of 0.01–70 Hz and sampled at 1000 Hz. Impedances were kept below 20 kΩ.

### 2.5. Eye movement recordings

The eye movements were recorded binocularly through a desktop-mounted Eyelink 1000 Plus eye tracking system (SR Research) at a sampling rate of 1000 Hz. The head position was stabilized via the chin rest of the tracker. A 9-point calibration was completed at the beginning of each paradigm. In addition, a 1-point drift correction check was performed at the beginning of each trial. Extra calibrations were performed whenever a fixation check failed. A calibration was accepted if the average validation error was below 0.5° and the maximum error across all points was below 1.0°.

### 2.6. Co-registration of Eye movements and EEG signal

Synchronization of eye movements and EEG was achieved by sending shared trigger pulses from the presentation PC (running Presentation, Neurobehavioral Systems Inc., Albany, CA) to the EEG and eye tracking computer on each trial through the parallel port. This allowed for accurate offline synchronization via the EYE-EEG extension for EEGLAB (http://www.eyetracking-eeg.org (Dimigen, 2020; Dimigen et al., 2011). After synchronization, the temporal offset between shared markers in both recordings rarely exceeded 1 ms.

### 2.7. Pre-processing of eye movement data

Three eye movement measures were used for data analysis, including first-fixation durations (FFD, the duration of the first fixation on a word), single fixation durations (SFD, fixation duration when a word only receives one first-pass fixation, and gaze durations (GD; the sum of fixations during the first pass reading of a word). The EyeLink Parser performed online eye-motion event detection and processing, including fixations and saccades, and recorded both raw gaze points and these results. Only fixations that occurred during the first-pass reading in correct trials were analyzed. Specifically, fixations on the area of interest were excluded, when the display change occurred too early or too late (i.e., when the display change took more than 10 ms before/after fixation onset on the target character). We also removed FFDs < 60 ms or > 600 ms (total number of excluded fixations: 1,844) and we excluded fixations on target characters in which participants blinked. In addition, we removed all trials with a wrong manual response to the animal question. And finally, we removed trials in which the display change was triggered already before saccade onset (‘early’ eye movements, *n* = 1,482, 25.72%, see discussion below), as preview characters were not presented in the parafoveal area. Taken together, we kept 5,103 observations for all participants.

### 2.8. EEG pre-processing

Offline, EEG data were digitally band-pass filtered using the *eegfiltnew.m* function of EEGLAB 2020.0 (Delorme & Makeig, 2004) toolbox for Matlab (version 2018b), between 0.1 Hz and 30 Hz (-6 dB/octave) and re-referenced to the average reference (Lehmann & Skrandies, 1980). Independent component analysis (ICA) was performed in order to identify the ocular artefacts using the EYE-EEG extension. ICA components that covaried with eye movements (variance threshold 1.1; Dimigen, 2020; Plöchl et al., 2012) were considered ocular artefacts and removed from the data. The corrected EEG signal was then segmented from 300 ms prior to and 700 ms after the first fixation onset on the target character and baseline-corrected by subtracting the voltages during the 150 ms interval preceding the fixation onset on the target word. Trials with amplitudes exceeding ±100 µV in any channel were automatically rejected from further analyses. ERPs were first averaged within and then across participants.

After eye movement and EEG preprocessing, we were left with a total of 3,393 observations for the target character in the boundary paradigm and 4,699 observations in the masked priming experiments. In each paradigm, participants had similar numbers of trials between repeated and unrelated conditions (boundary, *M* = 58.5, *SD* = 14.59, *range* = 31–81; masked priming, *M* = 81.02, *SD* = 4.27, *range* = 62–84). But the total trials left for the two paradigms was significantly different (*t* _(114)_ = −11.28, *p* < 0.001). The reason for this difference was that the boundary paradigm required more eye movements and therefore had more artefacts which could not be corrected compared with the masked priming paradigm (see Nikolaev et al., 2016) for discussion). Moreover, we excluded short fixations and “early” eye movements, which left fewer trials compared to the masked priming paradigm. Therefore, the two experiments were different in the degree of technical challenges, which resulted in a different amount of EEG trials that could eventually be included.

### 2.9. Data analysis

#### 2.9.1. Eye movement Statistical Analyses

Eye movement data were analysed with linear mixed-effects models within the R environment for statistical computing (R Core Team, 2015). We used the “lmer” function from the lme4 package (Bates et al., 2015) on log-transformed FFD, SFD and GD.

The within-subject factors of *Repetition* (identical vs. unrelated) and *Paradigm* (masked priming vs. boundary paradigm) were coded as fixed factors. Participants and items were specified as crossed random effects, with both random intercepts and random slopes (Barr et al., 2013). When we ran the models, we always began with full models that included the maximum random effects structure. But the slopes were removed if the model failed to converge (indicating over parametrization). The *p*-values were estimated using the “lmerTest” package, with the default Satterthwaites’s method for degrees of freedom and *t*-statistics (Kuznetsova et al., 2017).

#### 2.9.2. Pre-registered EEG/ FRP data analysis (https://osf.io/hk37q/)

##### Boundary paradigm

To investigate the neural correlates of the preview effects, we analysed FRP epochs time-locked to the fixation onset of the target. Based on previous studies, we selected 200–300 ms as the time window of preview positivity (Dimigen et al., 2012). Besides, as many studies observed the early N1 component, we selected 140–200 ms as the time window of N1. As the N1 effects and preview positivity have a scalp distribution on the occipital temporal regions with an average reference, we selected this area as regions of interest (ROI; left occipital-temporal area, LOT: PO9/PO7, and right occipital-temporal area, ROT: PO8/PO10) with a factor of Hemisphere (left vs. right). Therefore, Repeated Measure Analyses Variance (ANOVAs, with Bonferroni correction) were performed on the type of previews (*Repetition*, repeated vs. unrelated) and *Hemisphere* (left vs. right).

##### Masked priming paradigm

The data analyses of the masked priming paradigm were similar to the boundary paradigm, except for the selection of time windows. Based on the previous studies, the N1 effects occurred earlier than that of the boundary paradigm, therefore the time window of N1 in the masked priming paradigm was 120–175 ms. The N250 component had a similar time window as the preview positivity based on previous studies (Petit et al., 2006).

##### Comparison between two paradigms

To further investigate whether the repetition effects were similar across the two paradigms, we also performed three-way ANOVAs by assessing the interaction between the types of preview/prime and types of paradigms. Factors included *Repetition* (repeated vs. unrelated), *Paradigm* (masked priming vs. boundary paradigm) and *Hemisphere* (left vs. right). The Greenhouse-Geisser correction was applied to all effects with more than one degree of freedom in the numerator.

#### 2.9.3. Behavioural data analysis

Since manual responses were only executed following a 2000 ms blank screen interval, we did not analyze manual RTs in the study. In contrast, we calculated d-prime separately for probes in the preview/prime position and in the target position. This allowed us to quantify the visibility of the prime (masked priming) and the parafoveal preview (boundary paradigm) in the respective paradigm. In the following, we will call this ability to identify the prime/preview word *prime visibility*. In addition, we assessed the display change awareness for both paradigms.

## 3. Results

### 3.1. EEG results

#### 3.1.1. Traditional ANOVA results (proposed by preregistration)

Figure 2 presents the electrophysiological results obtained both the boundary paradigm and masked priming paradigm. To test whether repetition effects can be obtained for each paradigm, we first ran two-way ANOVAs for each time window of interest on the within-subject factors *Repetition* and *Hemisphere*. This was done for both the N1 and N250 components. Afterwards, three-way ANOVAs on *Paradigm*, *Repetition* and *Hemisphere* were performed to test whether or not repetition effects differ between paradigms in the two EEG components.

**Figure 2.**
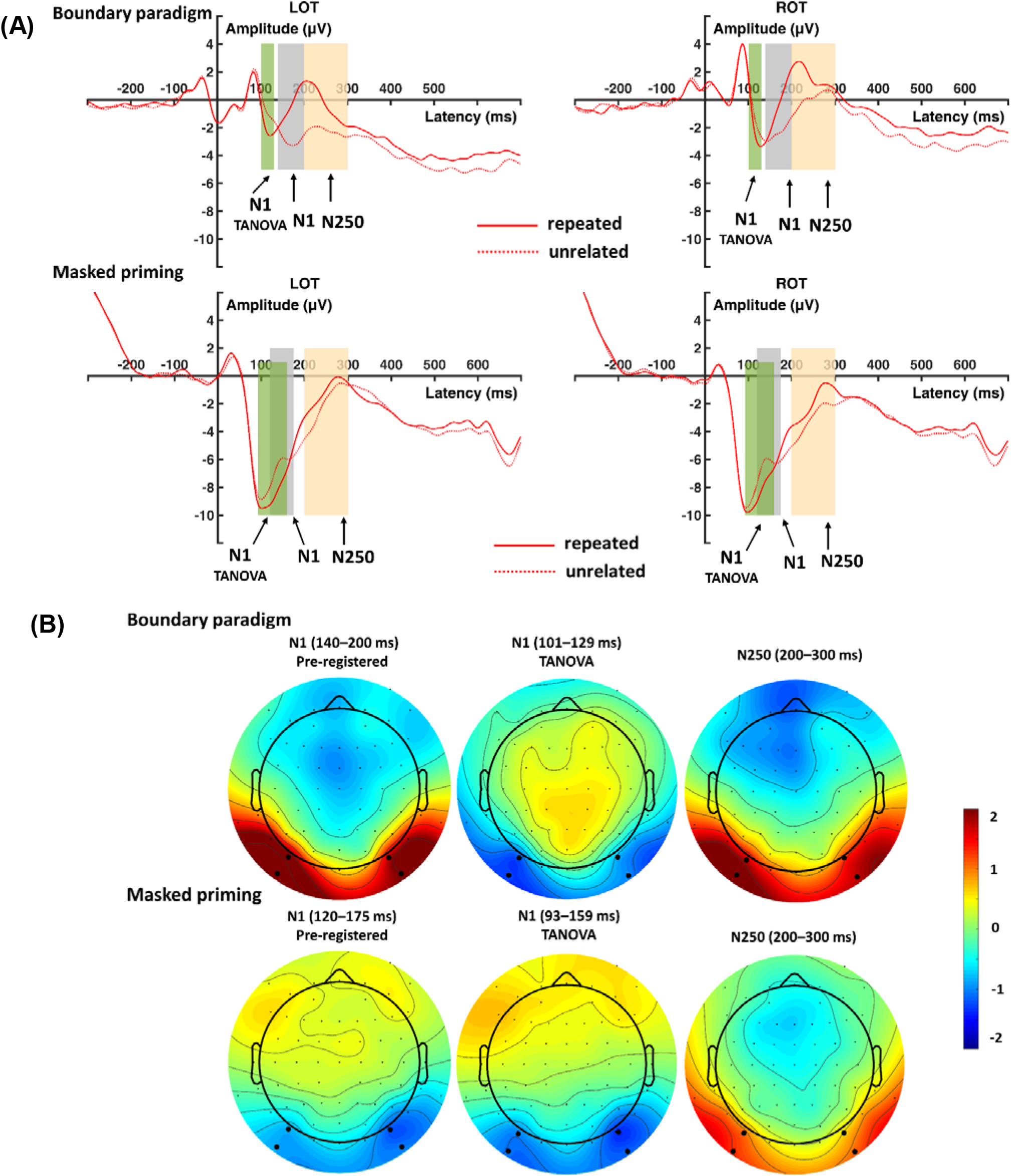
(A) Waveforms at left occipital-temporal (LOT) and right occipital-temporal (ROT) regions for each paradigm. Shading indicates the time windows used for the N1 and N250 components. (B) Effect topographies (repeated minus unrelated) of the N1 and N250 repetition effects for each paradigm. Black dots highlight the electrodes used to define the regions of interest (LOT and ROT).

##### N1 (pre-registered time window)

In the time-window of the N1 component which we had initially pre-registered for our analysis, we found that both the boundary paradigm and masked priming paradigm showed repetition effects (*boundary*: *F* _(1,28)_ = 41.03, *p* < 0.001, 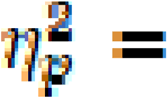 0.59; *masked priming*: *F* (1,28) = 34.41, *p* < 0.001, 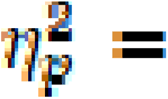 0.55, see Figure 2). However, the repetition effects in the two paradigms were in the opposite directions (increased for repeated characters in the masked priming paradigm and reduced for repeated characters in the boundary paradigm compared to unrelated characters). The other main effects and interactions were not significant (*F*s < 0.06, *p*s > 0.81).

We then ran the three-way ANOVAs on *Paradigm*, *Repetition* and *Hemisphere*. As anticipated from the reverse polarity of the effect, results showed that the repetition effects differed between the boundary paradigm and the masked priming paradigm (*Paradigm* × *Repetition*, *F* _(1,28)_ = 70.83, *p* < 0.001, 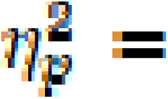 0.72). In addition, masked priming led to a larger, more negative N1 in response to the target than the boundary paradigm, regardless of repetition (*Paradigm*, *F* (1,28) = 46.33, *p* < 0.001, 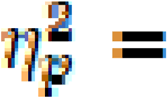 0.62), and the right hemisphere showed larger repetition effects than the left hemisphere (*Repetition* × *Hemisphere*, *F*(1,28) = 12.20, *p* = 0.002, 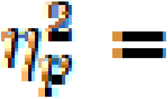 0.304; left vs. right: −3.83 vs. −4.02 μV). No other interactions were significant (*F*s < 0.005, *p*s > 0.94).

##### N250 (pre-registered)

In the N250 component, similarly, we ran a two-way ANOVA on *Repetition* and *Hemisphere* for each paradigm separately. Results showed that both the boundary paradigm and masked priming paradigm generated a reduced N250 negativity for repeated characters as compared to unrelated characters (*Repetition*, boundary, *F* _(1,28)_ = 36.62, *p* < 0.001, 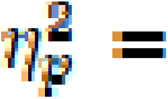 0.57; masked priming, *F*_(1,28)_ = 23.36, *p* < 0.001, 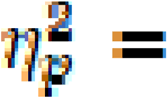 0.46). Furthermore, the left hemisphere showed a larger negativity than the right hemisphere, *F* _(1,28)_ = 9.91, *p* = 0.004, 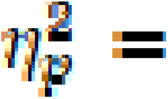0.26 (left vs. right: −1.23 vs. 0.87 μV). In addition, we observed a tendency that in the boundary paradigm, N250 repetition effects were numerically slightly larger (but not significantly so) in the right as compared to the left hemisphere (*Repetition* × *Hemisphere*, *F* _(1,28)_ = 3.00, *p* = 0.094, 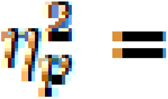 0.097). No other significant main effects and interactions were found (*F*s < 2.05, *p*s > 0.16).

We then ran a three-way ANOVA on *Paradigm*, *Repetition* and *Hemisphere*. The results showed that the masked priming paradigm led to a larger N250 negativity than the boundary paradigm (*Paradigm*: *F* _(1,28)_ = 6.58, *p* = 0.016, 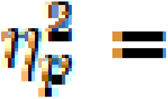 0.19), and the N250 was reduced for repeated as compared to unrelated targets (*Repetition*: *F*_(1,28)_ = 41.24, *p* < 0.001, 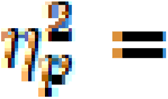 0.60), Similarly, the left hemisphere showed larger negativity than the right hemisphere (*Hemisphere*: *F*_(1,28)_ = 7.49, *p* = 0.011, 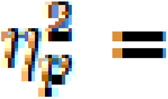 0.21). Interestingly, repetition effects were significantly larger in the boundary paradigm than that in masked priming (*Paradigm* × *Repetition*: *F* _(1,28)_ = 8.82, *p* = 0.006, 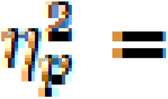 0.24; boundary vs. masked priming: −1.81 vs. −1.01 μV). The left hemisphere showed larger repetition effects than the right hemisphere (*Repetition* × *Hemisphere*: *F* _(1,28)_ = 4.52, *p* = 0.043, 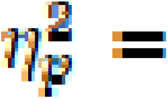 0.14). No other significant main effects and interactions were found (*F*s < 0.22, *p*s > 0.64).

#### 3.1.2. TANOVA results

As proposed in the pre-registration, we also explored the results with sample-by-sample Topographic Analyses of Variance (TANOVA; Koenig et al., 2011) on non-normalized (raw) maps comparing target ERPs/FRPs following valid primes/previews to those following invalid primes for unchanged conditions. The TANOVA were corrected for multiple comparisons through Global Duration Statistics (Koenig et al., 2011). Based on the TANOVA results, we selected the time windows in which repetition effects were significant (*p* < 0.05). As we were mainly interested in repetition effects corresponding to the N1 and N250 effects, we focused on effects occurring within 300 ms after the target was presented/foveated. TANOVA comparing repeated and unrelated targets identified two significant time windows each paradigm within 300 ms.

Results are shown in Figure 3. The N1 time window identified in the boundary paradigm, it was 101–129 ms. In the masked priming paradigm was 93–159 ms. Following the N1, the TANOVA identified a long significant time window in both paradigms, which may reflect an overlap of the N250 and N400 components. This time window was identified from 150 to 427 ms for the boundary paradigm, and from 176 to 453 ms for the masked priming paradigm. To further describe the time course and duration of repetition effects in the two paradigms, we plotted additional topographic maps in consecutive 20 ms time windows after fixation/stimulus onset (from 100 to 600 ms; see Figure 4). Together with the TANOVA results and the topographies, we found that time windows from the preregistration for the N1 overlapped with the N250 effect in both paradigms. Therefore, we ran an additional ANOVA on the N1 component using the time windows identified by TANOVA.

**Figure 3.**
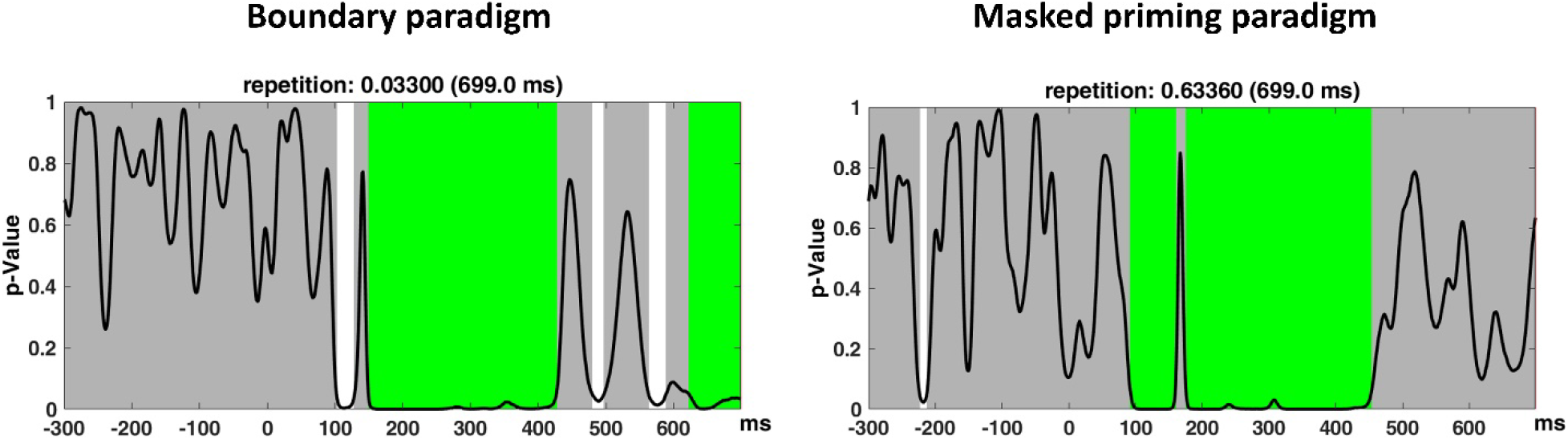
Sample-by-sample TANOVA for the boundary paradigm (left) and the masked priming paradigm (right) with global duration statistics. For the masked priming paradigm, the duration threshold was identified to be 51 ms, the threshold was then applied to the TANOVA plot, where significant periods longer than this estimated duration threshold are marked in green. For the boundary paradigm, the duration threshold was identified as 56 ms.

**Figure 4.**
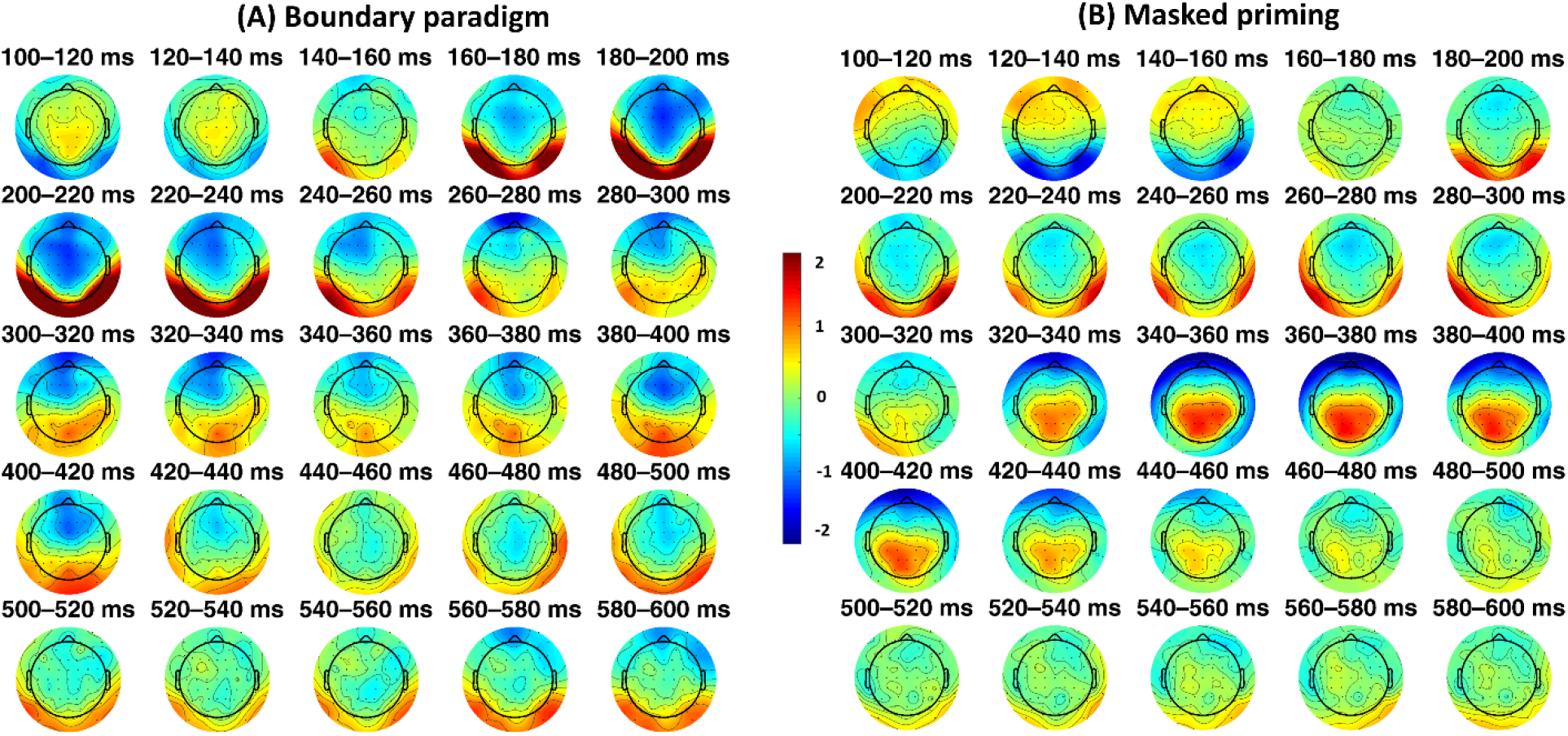
Scalp distribution of effects on the target character (repeated minus unrelated) averaged across consecutive 20 ms windows from 100 to 600 ms. Shown are results for the boundary paradigm (left panel) and the masked repetition priming paradigm (right panel).

The two-way ANOVA on each paradigm showed similar results in the identified N1 time windows (*boundary*, *F* _(1,28)_ = 14.15, *p* = 0.001, 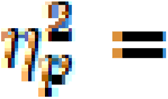 0.34; *masked priming*, *F* _(1,28)_ = 57.76, *p* < 0.001, 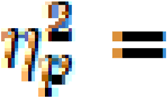 0.67), but the direction of the repetition effect in the boundary paradigm was reversed to the one in the pre-registered analysis (both paradigms showed increased repetition effects for repeated targets). No other main effects or interactions were significant (*F*s < 1.38, *p*s > 0.25).

The three-way ANOVA showed significant main effects of *Paradigm* (*F* _(1,28)_ = 74.59, *p* < 0.001, 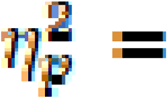 0.73) and *Repetition* (*F* _(1,28)_ = 38.99, *p* < 0.001, 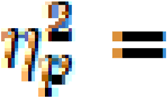 0.58). But the interaction between *Paradigm* and *Repetition* was not significant, *F* _(1,28)_ = 0.07, *p* = 0.79, 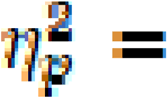 0.003. No other interactions were significant (*F* _(1,28)_ = 2.63, *p* = 0.12, 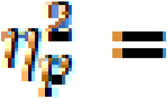 0.09).

### 3.2. Preview effect in fixation times (boundary paradigm)

The eye movement measures in the boundary paradigm did not show any preview effect, in contrast, at least in more later measures of fixation time (SFD, GD), participants tended to look at the target longer if the preview had been identical rather than unrelated (see Table 1). However, this tendency did not reach a significance in any of the eye movement measures (FFD: β = −0.004, SE =0.026, *t* = −1.48, *p* = 0.14; SFD: β = −0.019, *SE* = 0.031, *t* = −0.64, *p* = 0.52; GD: β = −0.01, *SE* = 0.02, *t* = −0.48, *p* = 0.63).

**Table 1.**
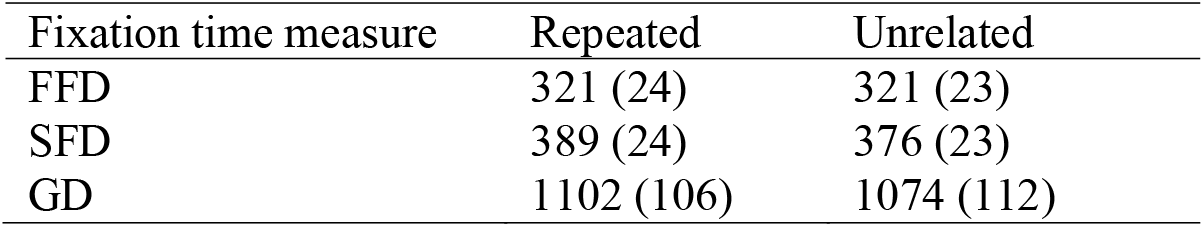
Means and standard errors of the fixation time measures (in milliseconds) for targets.

| Fixation time measure | Repeated | Unrelated |
| --- | --- | --- |
| FFD | 321 (24) | 321 (23) |
| SFD | 389 (24) | 376 (23) |
| GD | 1102 (106) | 1074 (112) |

### 3.3. Animal detection task and prime/preview visibility

Participants performed well in the animal task. In the boundary paradigm, they detected an average of 96.98% (*d*’ = 3.52) of probes in the preview position and 98.56% (*d*’ = 4.16) in the target position. In the masked priming paradigm, the mean accuracy of animal probes in the prime position was 96.94% (*d*’ = 3.55) and 98.95% (*d*’ = 4.33) in the target position.

The d-prime value was calculated from the proportion of hits on trials with animal names in the prime/preview position and false alarms on non-animal prime/preview trials. When the animal probes were in the target position, d-prime was higher than when the probes were in the preview/prime position (*F* _(1,28)_ = 57.09, *p* < 0.001, 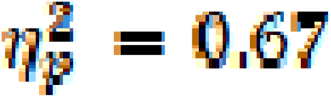). Neither the main effects of *Paradigm* (*F* _(1,28)_ = 0.96, *p* = 0.34) nor the interaction of paradigms and animal positions (*F* _(1,28)_ = 2.35, *p* = 0.13) was significant, suggesting the prime/preview visibility was similar across the two paradigms.

To test whether repetition effects were influenced by the prime’s visibility, we also correlated the d-prime with repetition effects in the N1 and N250 components for the two paradigms. We found that the d-prime in the preview/prime position was highly correlated with the one in the target position within a paradigm (masked priming: *r* = 0.45, *p* = 0.014; boundary paradigm, *r* = 0.632, *p* < 0.001). The N1 preview effects in the boundary paradigm were correlated with the d-primes in the preview and target position, but the other EEG repetition effects were not correlated with the d-primes (see Table 2 below). However, it should be noted that all these significant correlations were not significant after Bonferroni correction except the one for the *d*-prime in the preview position and target position of the boundary paradigm.

**Table 2.**
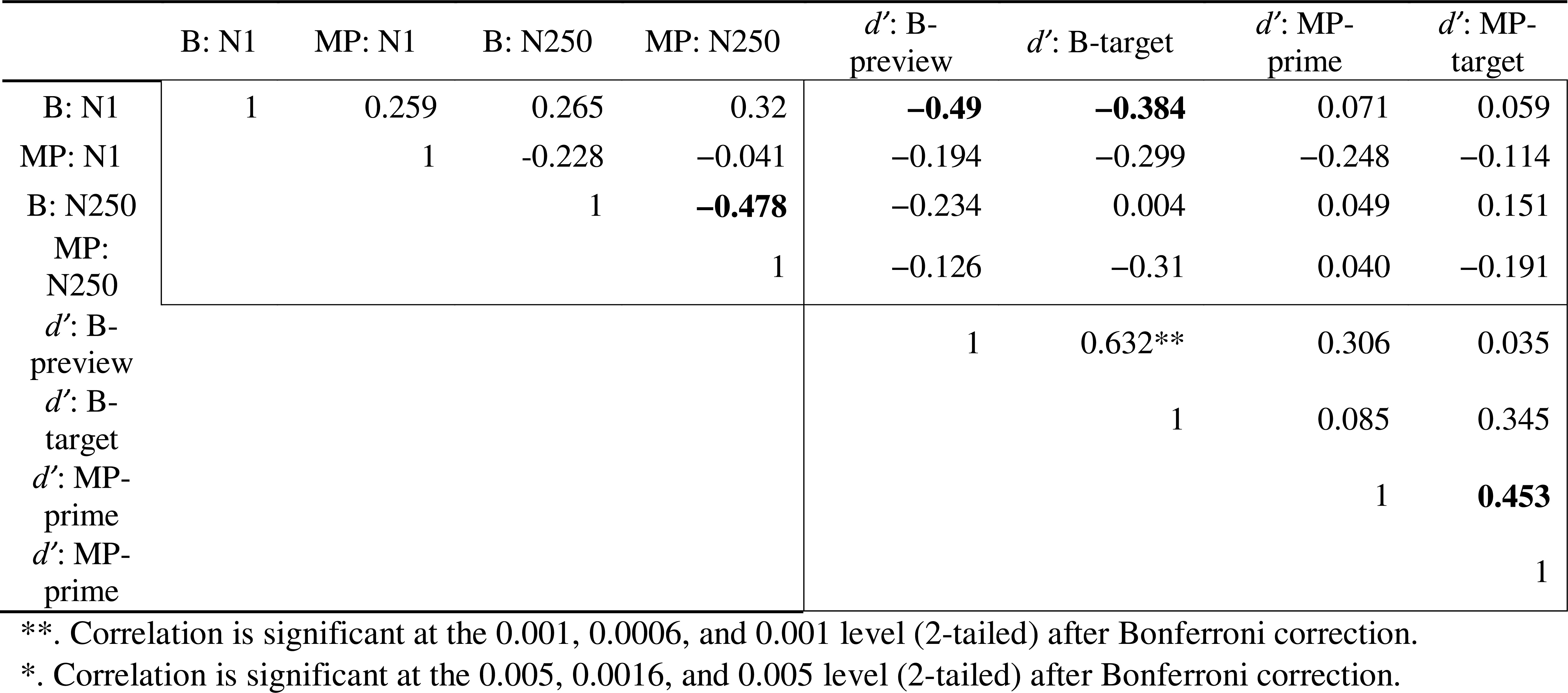
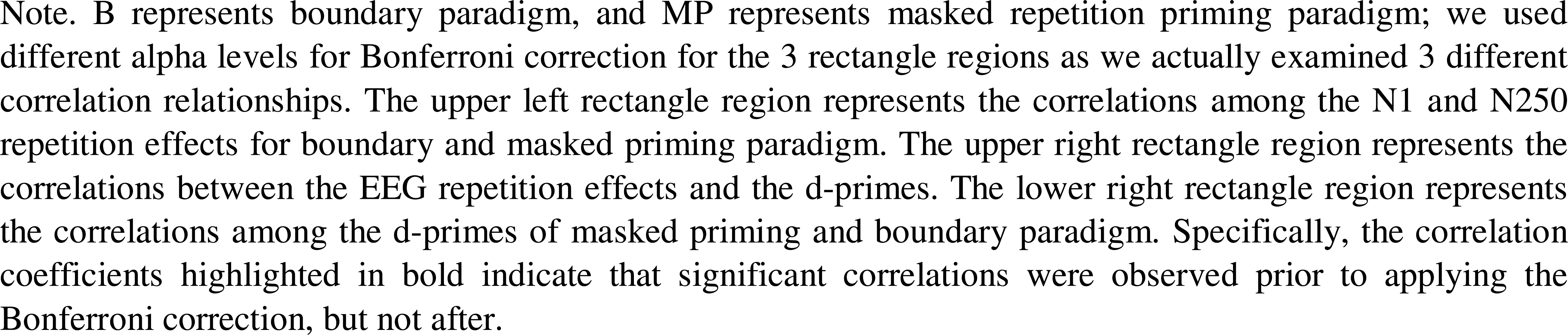
Correlations between repetition effects in the N1 and N250 components and the d-prime when animal characters were in prime/preview position or target position.

### 3.4. Display change awareness

Our measure of display change awareness assessed whether or not participants noticed that the display was manipulated. As expected, the display change awareness results showed that almost all participants were aware of the change of preview/prime, except 3 participants reported they did not note any change in the masked priming experiment. They reported an average of 29 (boundary, *M* = 29.38, *SD* = 21.8; masked priming, *M* = 29.48, *SD* = 23.64) trials showed changes (actually there were 200 changes in each paradigm, including 100 trials that involved only a font change but the same word). In addition, participants reported more frequent changes occurred in the masked priming paradigm than in the boundary paradigm, as they reported a larger percentage of changed trials in the masked priming paradigm than in the boundary paradigm (58.04% vs. 64.48%). This is likely explained by the inclusion of both a forwards and a backward mask making the screen changes more frequent.

## 4. Discussion

The current study directly compared the neural correlates underlying the effects of masked priming and parafoveal preview by presenting the same Chinese single character to the same participants in both paradigms. In each case, previews/primes were either identical or unrelated to the target characters. The neural correlates were measured by means of co-registering EEG and eye movements. We discussed the main findings below, beginning with the N250, followed by the N1 component.

### 4.1. N250 repetition effect

In the time window from 200–300 ms, we observed repetition effects in both the boundary paradigm and masked priming paradigm. Although the effect was larger in the boundary paradigm, the scalp distribution (occipital-temporal regions) and time course (200–300 ms) were similar in both paradigms. The preview positivity in the boundary paradigm has been suggested to reflect the preview-based facilitation of early stages in visual word recognition at visual feature and/or orthographic levels (Niefind & Dimigen, 2016). Consistent with this hypothesis, most eye movement researchers have located preview benefits in fixation durations primarily at the orthographic level (and to a lesser degree also at the phonological level) (Schotter et al., 2012b; Vasilev & Angele, 2017b). Also, in some EEG studies on orthographic processing, a separable N250 component was identified in addition to the N1, whereas in other studies, the N1 showed a decreasing flank that also covered the N250 time range. In addition, the time point-wise TANOVA revealed that the preview positivity starts around 150 ms after fixation onset. This result is consistent with the idea that saccade preparation in reading was estimated to take about 150–175 ms (Rayner et al., 1983). Thus, the temporal finding of the preview positivity is a plausible correlate of the facilitating brain processes that lead to the preview benefit in behaviour.

The N250 repetition effects in the masked priming paradigm has been suggested to reflect a process at the interface between sub-lexical and whole word representations and is sensitive to the degree of prime-target orthographic overlap; both partial repetitions and pseudoword repetitions benefit from the orthographic overlap between prime and target (for a review, see Grainger & Holcomb, 2010). The repetition effects in the boundary paradigm have been suggested to reflect a form of partial, trans-saccadic repetition priming that activates abstract orthographic and phonological representations (Dimigen et al., 2012). Given previous interpretations of these effects, it is possible that the preview positivity and the N250 repetition effects reflect a similar process in visual word recognition in the time window of 200–300ms.

However, as mentioned above, we found evidence that the N250 repetition effect is larger in the boundary paradigm than in masked priming. This phenomenon is consistent with the findings of Kornrumpf et al. (2016) who compared ERPs in an RSVP-with-flanker paradigm (i.e., RSVP presentation with parafoveal flanker words providing previews) and FRPs in the boundary paradigm and found that the preview positivity (200–280 ms) was substantially smaller in ERPs (without eye movements) than in FRPs. This may be due to the mandatory shift of visuo-spatial attention toward *n* + 1 occurring in natural reading that is coupled to the preparation of the saccade. In contrast, such an attention shift is likely unnecessary in the RSVP-flanker paradigms which ask participants to maintain constant eye fixation while words are presented one by one in foveal vision at a fixed pace (e.g., masked priming paradigm). Therefore, we think that the basic processes reflected by the N250 masked priming effect and the preview positivity may be very similar, but that they are more pronounced in the latter paradigm, possibly because of deep level of consciousness of the stimuli, or specific word parts that readers process (e.g., word centre vs. initial letters), or duration of the previews/primes.

Although many researchers agree that the N250 effects between 200–280 ms reflects preview benefits, it is under debate that the N250 effects may be also the consequence of preview costs, as readers may generate implicit expectations about the upcoming words. How to explain the effect is actually determined by the baseline used for computing the preview effects (e.g., Yan et al., 2012). However, this argument can be viewed as two sides of a coin within the predictive coding framework (e.g., Summerfield et al., 2008). The framework of predictive coding (Friston, 2005) assumes that humans constantly generate predictions about likely sensory inputs, whereas the actual sensory input serves as an error signal to allocate attention to unpredicted events. Within this framework, it is possible that expectations before the saccade may be violated after the saccade following invalid previews (as suggested by Kornrumpf et al., 2016). Furthermore, some studies have found that the N250 modulations showed some resemblance to the visual MMN (Stefanics et al., 2014), and therefore may reflect a mismatch signal (Rao & Ballard, 1999). When the upcoming words fulfil the expectations (i.e., identical previews/primes), the processing is facilitated.

### 4.2. N1 repetition effect

In the masked priming paradigm, we obtained a typical repetition effect in the N1 component (e.g., Petit et al., 2006), where targets elicited larger amplitudes after identical than after unrelated primes at occipital-temporal electrodes. While the timing of this effect coincides with a broad negativity in the ERP with several small peaks, the early time window of this effect around 100 to 150 ms may correspond to the P1-N1 transition in regular ERPs, where the P1 typically occurs around 100 ms, and the N1 at around 150 ms (e.g., Grainger & Holcomb, 2009). The repetition priming effect in this time window therefore would also be in agreement with a forward-shift of the N1 component latency, although this might not be visible in masked priming experiments due to the overlap with ERP components elicited by the preceding mask and prime.

The TANOVA results revealed that the N1 repetition effects in the boundary paradigm occurred earlier (i.e., 101–129 ms) than the time window proposed in our pre-registration (140– 200 ms). This earlier time window identified by the TANOVA may be due to that different approach for identifying time windows. In the pre-registration, we selected the time window of the N1 repetition effect based on previous studies (Dimigen et al., 2012), but the data-driven approach can capture the effect more precisely.

The early N1 effect consisted in a larger negativity after a valid than an invalid preview at occipital-temporal electrodes. Previous studies which have reported early N1 preview effects (e.g., Niefind & Dimigen, 2016), in which they found an early identity preview effect with a larger negativity after unrelated previews than valid previews. This pattern was consistent with the typical preview positivity in the later time window (200–300ms). Therefore, these previously found N1 preview effects between 160 and 200 ms should be considered as early part of preview positivity. This hypothesis was supported by the TANOVA results, indicating a preview positivity starting occurred as early as 153 ms after fixation onset. These findings provide further insights into the mechanisms underlying the classic preview effect. This early N1-like preview effect, similar to the N1 repetition effect in the masked priming paradigm, may reflect the feature overlap processing between prime and target. When targets and previews/primes are identical, the targets have a larger number of visual features shared with the primes or previews as compared to mismatched pairs, and targets after identical primes/previews have a processing benefit. Therefore, we think the N1 effect in the masked priming and boundary paradigm actually reflect the same mechanism perceptual priming at the visual feature level.

In addition, we think that the pattern of an N1 increase and a N250 decrease (e.g., also qualitatively seen in Dimigen & Ehinger, 2021, see their Figure 8) could potentially occur due to a latency shift of the N1 after valid previews or primes. However, in this case, the size of the N1 effect should correspond to the size of the N250 effect, which was not the case in the current study. So, a potential latency shift could only partially explain the N250 effect. Therefore, one speculation is that the facilitation effect does not just seem to be the result of speeding up N1 processing, but presumably it also consists of reduced activation during the N250 time range or potentially acceleration of subsequent processing.

We did not focus on the N400 component in the current analyses. However, we did replicate an N400 attenuation by repeated previews/primes in both paradigms (see topographies in Figure 4; see also time windows identified in TANOVA). The presence of an expected N400 effect provides further evidence that the early effects we obtained were typical.

### 4.3. No behaviour preview/priming facilitation effects

We did not find the typical preview effects in fixation times. To the contrary, readers showed a trend towards shorter fixations after unrelated than identical previews, at least in later measures of fixation time (SFD, GD). This numerical trend, although not significant, was therefore reversed as compared to the typical pattern in eye-tracking studies during sentence reading (e.g., Schotter, 2013; Yang et al., 2012). Likely, in our simplified reading paradigm, there was no more upcoming visual information to the right of the target word, which further increased fixation durations beyond that in natural reading situations (e.g., Rayner et al., 2006). This suggests that for the emergence of preview effects, a more natural reading setting with ongoing saccade behaviour is needed.

### 4.4. Effects of prime/preview visibility on ERP measures

In the masked priming paradigm, neither the early N1 nor the N250 repetition effect were influenced by the prime’s visibility – i.e., whether the participant could correctly classify whether this briefly presented and masked word was an animal or not; in contrast, in the boundary paradigm, preview visibility seems to influence the N1 preview effect, but not the later N250 effect. A previous masked priming study (Holcomb et al., 2005) found that prime visibility in a masked priming paradigm only influenced the N400 but not earlier effects (e.g., N250), suggesting that prime visibility may have little influence on early stages of visual word recognition for the target word. In a masked priming paradigm, where readers fixate on the centre of the screen, the primes are effectively masked to the greatest degree. In contrast, in a boundary paradigm, readers are allowed to move their eyes freely. In this case, when the participants’ task is to judge whether or not there was an animal probe in a trial, they may develop a strategy to move their eyes to the target word later and use their extrafoveal vision to try to identify the preview. This hypothesis is supported by the more robust correlation between the N1 preview effect and sensitivity (d-prime) when the animal probe was in the preview position than the one when the animal probe was in the target position. This may explain why the preview visibility only influences the repetition effects in the boundary paradigm but not in the masked repetition priming paradigm. In addition, the N1 component is sensitive to the degree of overlap between the target and prime items while the N250 component is more sensitive to orthographic information. This may explain why preview visibility influenced only the early N1 preview effect, but not the later component.

Another interesting finding from the correlation analysis was that, across participants, the size of the N250 preview effect in the boundary paradigm correlated with the size of the N250 repetition effect in the masked priming paradigm. The results provide further support that these two effects have a similar neural basis; this holds true even though the N250 effect was larger in the boundary paradigm than in the masked priming paradigm.

### 4.5. Limitations and future directions

We are aware of some limitations of the current study. First, we noted that the number of remaining trials in the boundary paradigm was smaller than in the masked priming paradigm, which means that our analysis might be less sensitive to pick up effects in the boundary paradigm One reason is that there were generally more EMG artefacts in the boundary paradigm. Another reason for the larger loss of trials were premature saccades in some trials, that occurred before the preview was even presented. These early eye movements may have resulted from the rather long fixation check at the beginning of each trial. However, even though fewer trials were left for analysis of the boundary paradigm compared to the masked priming paradigm, we still observed a larger N250 repetition effect in the boundary paradigm than in the masked priming paradigm, suggesting that the effect is robust.

Second, as mentioned earlier, we did not find a preview effect in fixation times in our less natural reading setting. We chose this single-saccade version of the boundary paradigm in order to make the two paradigms directly comparable. Future studies may consider an experimental design closer to a natural reading situation, for example, by embedding the words into sentences or word lists, or by adding another fixation point to the right of the targets, so there is also an outgoing saccade from the target word.

In a masked repetition priming paradigm, the prime is backward-masked after a brief foveal presentation, which renders it invisible. In contrast, in the boundary paradigm, the preview is presented parafoveally but usually with a longer duration (corresponding to the gaze duration on the fixation cross). Although N250 effects differed in size, the current results suggest that the consequences of these two rather different manipulations are qualitatively surprisingly similar at the neural level. Future studies could address in more detail the question to what degree a brief backward-masked prime is comparable to a longer but parafoveally degraded and visually crowded preview and at which level the previews or primes are processed. This could be done by introducing new experimental conditions, for example by shifting the prime into parafoveal vision in the masked priming paradigm (Pernet et al., 2007) or by masking the preview in the boundary paradigm (cf., Hohenstein et al., 2010; Rayner et al., 2006).

## Conclusion

In sum, qualitatively similar effects on the N1 and N250 components were obtained in the boundary and masked priming paradigm. The similar scalp distributions, time courses and the directions of effects indicate that the neural mechanisms underlying the two paradigms are similar or strongly overlapping. Furthermore, the boundary paradigm, which allows readers to execute saccades had a bigger impact on the N250 component as compared to the very early stage of visual word recognition, possibly because readers are more actively engaged in reading.

## Supporting information

Supplementary experiment

## Acknowledgements

This work was supported by the General Research Fund of the Research Grants Council of Hong Kong (RGC-GRF 14616418), as well as by the Germany/Hong Kong Joint Research Scheme of the Research Grants Council of Hong Kong (G-CUHK409/18) and the German Academic Exchange Service (DAAD, Project 57447990).

## Data availability

All data and codes have been made publicly available at https://osf.io/hk37q/.

## Footnotes

1 Note that we accidently set the duration of the target presentation to 1000 ms and therefore longer than the target duration in the masked priming paradigm. This is likely not crucial, as we exclusively analyze effects that occur well before 500 ms. However, to further test whether the duration of target had any substantive influence on the early electrophysiological effects, we ran a supplementary study in which the duration of targets was set to 500 ms in both paradigms. This is reported as a supplementary experiment in the Appendix).

