## Supplementary experiment for "Neural mechanism underlying preview effects and masked priming effects in visual word processing"

During the main experiment, the targets in the boundary paradigm were accidently presented for 1000 ms rather than 500 ms (as in the masked priming paradigm). Because all ERP components we focused occurred before 500 ms, we think this would not influence the results much. Nevertheless, to make the two paradigms even more comparable, we ran an additional experiment in which we set the duration of the target presentation in both paradigms to 500 ms.

### Participants

An additional 16 native Cantonese Chinese participants (7 females; mean age = 20.55 years, range = 18–22 years) were tested in both experiments. Recruitment criteria for participants were the same as in the proper experiment. Data from three subjects were excluded from the analysis due to a small number of remaining trials (*n* < 15).

### Materials and procedure

All materials and procedures were identical to the main experiment, except that the duration of the target presentation in the boundary paradigm (relative to the eyes triggering the invisible boundary) was set to 500 ms.

### Results

#### ***Traditional ANOVA results (proposed by preregistration)***

We took the same strategy in the formal experiment to analyze the data from this supplementary experiment. We first run two-way ANOVAs for each time window of interest with within-subject factors *Repetition* (unrelated vs. repeated), and *Hemisphere* (left vs. right).

**N1.** In the N1 component, we found that both the boundary paradigm and masked priming paradigm found repetition effects (*boundary*, *F* _(1,12)_ = 12.90, *p* = 0.001, 0.52; *masked* *priming*, *F* _(1,12)_ = 20.99, *p* = 0.001, 0.64). Similar to the formal experiment, we observed that the repetition effects between the two paradigms were opposite (increased for repeated characters in the masked priming paradigm and reduced for repeated characters in the boundary paradigm compared to unrelated characters). In addition, the repetition effect was slightly larger in the right hemisphere compared to the left hemisphere (*Repetition* × *Hemisphere*, *F* _(1,12)_ = 3.84, *p* = 0.074, 0.24). The other main effects and interactions were not significant (*F*s < 1.51, *p* > 0.24).

We then ran the three-way ANOVAs on *Paradigm*, *Repetition* and *Hemisphere*, results showed repetition effects were larger in the boundary paradigm than that in the masked priming paradigm (*Paradigm* × *Repetition*, *F* _(1,12)_ = 30.75, *p* < 0.001, $?_{p\text{?}}^{2}=$ 0.72). In addition, masked priming led to larger N1 negativity in the subsequent targets than boundary paradigm (*Paradigm*, *F* _(1,12)_ = 34.94, *p* < 0.001, 0.74), and the right hemisphere showed slightly larger activation for repetition effects than the left hemisphere (*Repetition* × *Hemisphere*, *F* _(1,12)_ = 3.28, *p* = 0.095, 0.215; left vs. right: –3.32 vs. –3.82 µV). No other interactions were significant (*F*s < 2.16, *p* > 0.17).

**N250.** In the N250 component, similarly, we ran a two-way ANOVA on *Repetition* and *Hemisphere* for each paradigm, results showed that both the boundary paradigm and masked priming paradigm found reduced N250 negativity for repeated characters compared to unrelated characters (*Repetition*, boundary, *F* _(1,12)_ = 4.17, *p* = 0.064, 0.26; masked priming, *F* _(1,12)_ = 3.23, *p* = 0.097, 0.21). In addition, we observed that in the masked priming paradigm, the repetition effects were slightly larger in the right hemisphere than left hemisphere with a marginally significant interaction between *Repetition* and *Hemisphere* (*F* _(1,12)_ = 3.30, *p* = 0.095, 0.22). No other significant main effects and interactions were found (*F*s < 1.68, *p* > 0.22).

We then ran the three-way ANOVA on *Paradigm*, *Repetition* and *Hemisphere*, results showed a masked priming paradigm led to larger N250 negativity in the subsequent targets than the boundary paradigm(*Paradigm*, *F* _(1,12)_ = 18.80, *p* = .001, 0.61), and the N250 was reduced for repeated compared to unrelated targets (*Repetition*, *F* _(1,12)_ = 4.96, *p* = 0.048, 0.29),

The left hemisphere showed larger activation for repetition effects than the right hemisphere, *Repetition* × *Hemisphere*, *F* _(1,12)_ = 3.95, *p* = 0.07, 0.25. No other significant main effects and interactions were found (*F*s < 0.39, *p*s > 0.55).

#### ***TANOVA results***

Similar to the formal experiment, we ran a time-point-wise TANOVA comparing repeated and unrelated targets for each paradigm. For the masked priming paradigm, TANOVA identified 4 time windows after stimulus onset, which are 113–152 ms (N1), 194–215 ms (N250), 316–457 ms and 606–642 ms. For the boundary paradigm, similarly, 4 time windows were identified, which are 103–129 ms (N1), 162–217 ms (N250), 325–349 ms, and 369–386 ms (see Supplementary Figure 1A).

As TANOVA results revealed, the N250 repetition effects were narrowed in a shorter time window for both paradigms and no longer overlapped with the time window of the N400 effect. Therefore, we further tested the N1 and N250 effects with the time windows identified by TANOVA.

**N1.** Similar to the traditional analyses, the two-way ANOVA on *Repetition* and *Laterality* for each paradigm, results showed that both the boundary paradigm and masked priming paradigm found increased N1 negativity for repeated characters compared to unrelated characters (*Repetition*, boundary, *F* _(1,12)_ = 12.25, *p* = 0.004, 0.51; masked priming, *F* _(1,12)_ = 53.55, *p* < 0.001, 0.16). In addition, the right hemisphere showed larger activation than the left hemisphere for the repetition effects in the boundary paradigm (*Hemisphere*, *F* _(1,12)_ = 6.25, *p* = 0.028, 0.34).

The there-way ANOVA on *Paradigm*, *Repetition* and *Hemisphere* revealed no significant differences in the repetition effects between the two paradigms (*Repetition* × *Paradigm*, *F* _(1,12)_ = 2.76, *p* = 0.12, 0.19). And masked priming showed larger N250 negativity than boundary paradigm, *F* _(1,12)_ = 2.76, *p* = 0.12, 0.19. Repeated targets elicited larger negativity than unrelated targets, *F* _(1,12)_ = 48.02, *p* < 0.001, 0.80. In addition, the boundary paradigm showed larger hemisphere differences (*Paradigm* × *Hemisphere*, *F* _(1,12)_ = 4.20, *p* = 0.063, 0.26). Most importantly, we observed a three-way interaction between *Repetition*, *Paradigm* and *Hemisphere*, showing that the repetition effect was larger in the boundary paradigm than masked priming paradigm with larger activation in the right hemisphere (*Paradigm* × *Hemisphere* × *Repetition*, *F* _(1,12)_ = 10.88, *p* = 0.006, 0.48). The other main effects or interactions were not significant (*F*s < 1.4, *p*s > 0.26).

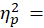

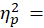

**N250.** The two-way ANOVA on *Repetition* and *Hemisphere* for each paradigm showed that both the boundary paradigm and masked priming paradigm found reduced N250 negativity for repeated characters compared to unrelated characters (*Repetition*, boundary, *F* _(1,12)_ = 22.90, *p* < 0.001, 0.66; masked priming, *F* _(1,12)_ = 11.25, *p* = 0.006, 0.48). In addition, the right hemisphere showed larger activation than the left hemisphere for the repetition effects in the boundary paradigm (*Hemisphere* × *Repetition*, *F* _(1,12)_ = 5.53, *p* = 0.037, 0.32). The other main effects or interactions were not significant (*F*s < 0.58, *p*s > 0.46).

The three-way ANOVA on *Paradigm*, *Repetition* and *Hemisphere* did not find any interactions, although the repetition effects on the boundary paradigm were larger compared to masked priming (*Paradigm* × *Repetition*, *F* _(1,12)_ = 0.47, *p* = 0.51, 0.04, mean amplitude: 1.99 vs 1.65). The repetition effect was significant (*F* _(1,12)_ = 24.28, *p* < 0.001, 0.67), with repeated targets eliciting reduced negativity compared to targets after unrelated primes/previews. And masked priming paradigm had larger amplitudes than the boundary paradigm (*Paradigm*, *F* _(1,12)_ = 17.09, *p* = 0.001, 0.59).

In summary, the new data set from a small group of 13 participants mainly replicated the findings of the main experiment, suggesting that early effects were not influenced in any major way by the different target duration. Although the interactions between repetition and paradigm did not reach significance in this small sample of thirteen participants, the results still showed the same consistent trend that the boundary paradigm had larger repetition effects than the masked priming paradigm. The nonsignificant interaction may result from the small sample size.

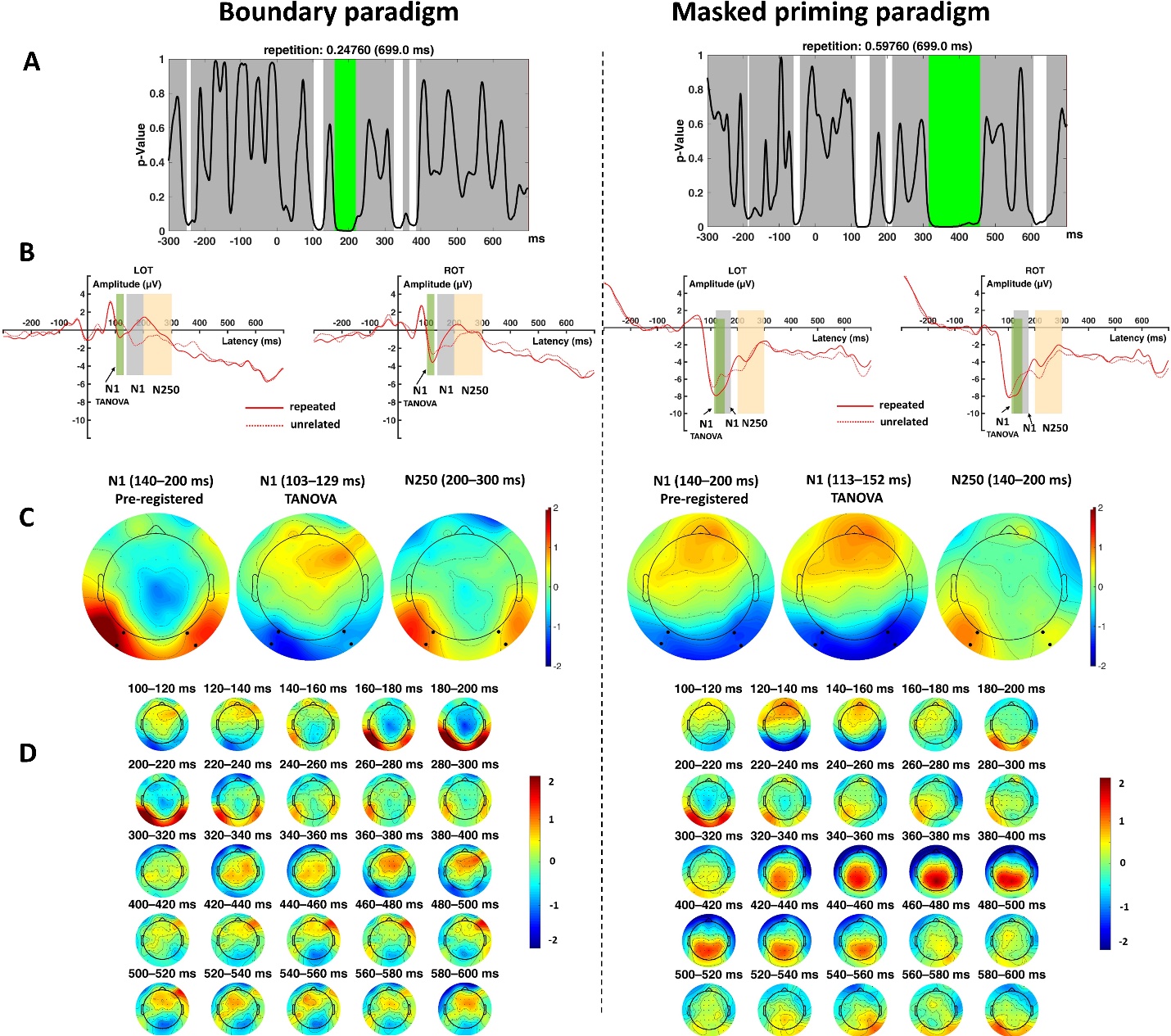

*Supplementary Figure 1.* Detailed results for the preview benefit (left panel) and masked priming repetition effect (right panel). (A) TANOVA results with global duration statistics correction. For the boundary paradigm, the duration threshold was identified as 45 ms, the threshold was then applied to the TANOVA plot, where periods longer than the estimated duration threshold are marked in green. For the masked priming paradigm, the duration threshold was identified as 450 ms. (B) Repetition effects (unrelated minus repeated) in left occipital–temporal (LOT, PO7 and PO9) regions and right occipital–temporal (ROT, PO8 and PO10) regions for boundary paradigm (left panel) and masked priming paradigm (right panel). (C) Topography maps of repetition effect (repeated minus unrelated) in N1 and N250 for each paradigm. (D) Temporal evolution of repetition effect (repeated minus unrelated) in successive 20 ms time windows between 0 and 600 ms after stimulus/fixation onset.
